# KRT7^CreER^ Ferrets Enable Lineage Tracing of Ductal Epithelia and Pancreatic Remodeling

**DOI:** 10.64898/2026.10.16.759748

**Authors:** Yaling Yi, Yu Sun, PeiPei Wu, zheqing hu, Yuhui Weng, Feng Yuan, Xiaoming Liu, Ziying Yan, Xingshen Sun, John F. Engelhardt

**Author notes:** Co-first author (Yaling Yi, Yu Sun). **Corresponding author**: Xingshen Sun, Pulmonary, Allergy, and Critical Care Medicine, Department of Medicine, University of Alabama at Birmingham, Birmingham, Alabama, 35294, USA, John F Engelhardt, Pulmonary, Allergy, and Critical Care Medicine, Department of Medicine, University of Alabama at Birmingham, Birmingham, Alabama, 35294, USA.

## Abstract

Keratin 7 (KRT7) is expressed in simple and ductal epithelia and is associated with epithelial plasticity during development, injury, and disease. However, the in vivo fate and regenerative behavior of KRT7⁺ cells remain poorly defined because lineage-tracing models are lacking in species that closely recapitulate human disease. To address this gap, we generated *KRT7*^CreER^ ferrets and crossed them with *ROSA^mTmG^* Cre-reporter ferrets, with or without the *CFTR^G551D/G551D^*mutation. Following tamoxifen induction, KRT7-derived EGFP⁺ cells were identified in glandular and ductal epithelia across multiple organs. Pulse-chase lineage tracing revealed that KRT7⁺BSND⁺ cells turn over rapidly during salivary gland development. In the pancreas, KRT7 lineage-labeled cells are present at all levels of the ductal tree under homeostatic conditions. In cystic fibrosis (CF) ferrets, KRT7-derived centroacinar cells and intercalated ductal lineages persisted and underwent marked remodeling, characterized by expansion of the SOX9⁺ centroacinar cell pod into intercalated ducts. Although acinar cells were not lineage-labeled in the healthy pancreas, *EGFP⁺/CELA3B*⁺ cells emerged in the CF pancreas and in acinar-derived organoids, consistent with induction of KRT7 during acinar-to-ductal metaplasia. These findings establish *KRT7^CreER^* ferrets as a versatile model for studying ductal epithelial lineage dynamics, pancreatic remodeling, and epithelial plasticity in both homeostatic and pathological conditions.

## Introduction

Keratins are a large family of intermediate filament proteins that form the structural backbone of epithelial cells, allowing epithelial tissues to withstand physiological stress and injury. In mammals, keratins are encoded by 54 evolutionarily conserved genes, including 28 type I (acidic) and 26 type II (basic to neutral) members (1, 2), and their expression is tightly controlled in a tissue-specific and developmentally dynamic way, with different epithelial cell types expressing unique combinations of keratins (3). This highly selective expression pattern provides a robust framework for defining epithelial identity and has been widely used in lineage-tracing techniques to study cell fate, plasticity, and regeneration in vivo.

Among these keratins, keratin 7 (KRT7; cytokeratin 7, CK7) has gained increasing attention due to its emerging roles in epithelial biology and disease (4). Under physiological conditions, KRT7 expression is associated with epithelial proliferation, migration, and epithelial–mesenchymal transition, which are essential for embryonic development, immune responses, and wound healing (4). Under pathological contexts, including cancer (4), injury (5, 6), stress (7), and tissue regeneration (4), KRT7 expression is dysregulated. In these disease states, KRT7 is often upregulated in response to epithelial stress and is strongly expressed in ductal reactions, metaplastic lesions, and pancreatic ductal adenocarcinoma. Higher KRT7 levels are associated with increased epithelial growth, migration, and cellular plasticity, as well as more aggressive tumor behavior and poor prognosis in pancreatic cancer (4). Overall, these findings highlight KRT7⁺ cells as active players in disease progression rather than passive structural components.

Despite growing recognition of KRT7’s role in epithelial pathology, much of our current understanding of its biology derives from mouse models (8, 9) and studies of pathogenic keratin mutations or aberrant expression in human diseases (10, 11). However, mouse models do not fully replicate several key aspects of human pancreatic pathology, especially cystic fibrosis (CF)–related pancreatic disease. In contrast, CF ferrets faithfully model not only CF-associated lung disease but also pancreatic disorders, including exocrine pancreatic insufficiency and CF-related diabetes (12, 13). The CF pancreas is characterized by impaired secretion of ductal fluid and bicarbonate, luminal obstruction, inflammation, and progressive tissue damage (14–16), features that closely mirror human CF pancreatic disease. Importantly, studies in CF ferrets have shown that ductal epithelial dysfunction is a primary initiating factor that triggers subsequent injury and repair responses. These studies also reveal that KRT7-expressing ductal cells are the exclusive source of CFTR expression in the pancreas (17–19).

Compared with mouse salivary glands, ferret major salivary glands appear more similar to human than to mouse salivary glands in several practical and anatomical respects, especially with respect to ductal organization, the absence of rodent-specific specializations, and the translational relevance of CFTR-dependent exocrine biology, although the direct comparative literature is limited(20, 21). In rodents, the submandibular gland contains a prominent granular convoluted tubule compartment between the intercalated and striated ducts, a specialization without a clear human counterpart (22–24). These species differences are important when interpreting ductal lineage relationships and epithelial specialization across model systems. In human salivary duct epithelia, CFTR contributes to chloride and bicarbonate transport and helps regulate the final composition of saliva(25–27). These processes are relevant to cystic fibrosis (CF), where altered salivary composition, reduced buffering capacity, xerostomia, and increased oral health complications have been reported in patients (28, 29).

Despite these advances, the in vivo fate, lineage stability, and injury-responsive behavior of specific ductal subsets, especially KRT7⁺ ductal cells in the salivary gland and pancreas, remain poorly understood. To address these gaps, we created a *KRT7*^CreER^ ferret model that allows for inducible genetic labeling of KRT7⁺ epithelial cells. When integrated with our established ROSA^mTmG^ Cre-reporter ferret and CF ferrets, this model demonstrates strong specificity and inducibility for tracking pancreatic ductal cells under both physiological and pathological conditions. Using pulse-chase experiments, we also identified a unique ductal lineage within the salivary gland. In addition to the pancreas and salivary gland, KRT7⁺ cells were lineage-traced in multiple additional organs. Collectively, these findings establish *KRT7*^CreER^ ferrets as a resource for dissecting epithelial cell lineage dynamics and function across a spectrum of diseases that are uniquely recapitulated in ferrets.

## Results

### Generation of *KRT7*^CreER^ ferrets using EASI-CRISPR

To selectively label KRT7-expressing glandular and ductal epithelia, we generated *KRT7*^CreER^ ferrets by inserting an *IRES-CreERT2* cassette into the 3′ UTR of the *KRT7* gene using EASI-CRISPR. A long ssDNA donor template and a guide RNA were designed to target the region downstream of the *KRT7* stop codon. The ssDNA donor cassette contained an internal ribosome entry site (IRES) followed by *CreERT2*, enabling independent translation of CreERT2 from the endogenous *KRT7* transcript. The cassette was flanked by ∼900 bp homology arms (Fig. 1A). The Cas9/sgRNA ribonucleoprotein complex and *KRT7-IRES-CreERT2* ssDNA donor were co-microinjected into ferret zygotes. Offspring carrying the targeted allele were identified by PCR using primers flanking the homology arms, confirming *CreERT2* integration at the *KRT7* locus (Fig. 1B). PCR-based genotyping indicated targeted insertion had occurred in 5 of the 13 kits born (4 survived for further screening). Correct integration was further validated by sequencing the expected 5′ and 3′ junctional PCR fragments (Fig. 1A).

**Figure 1.**
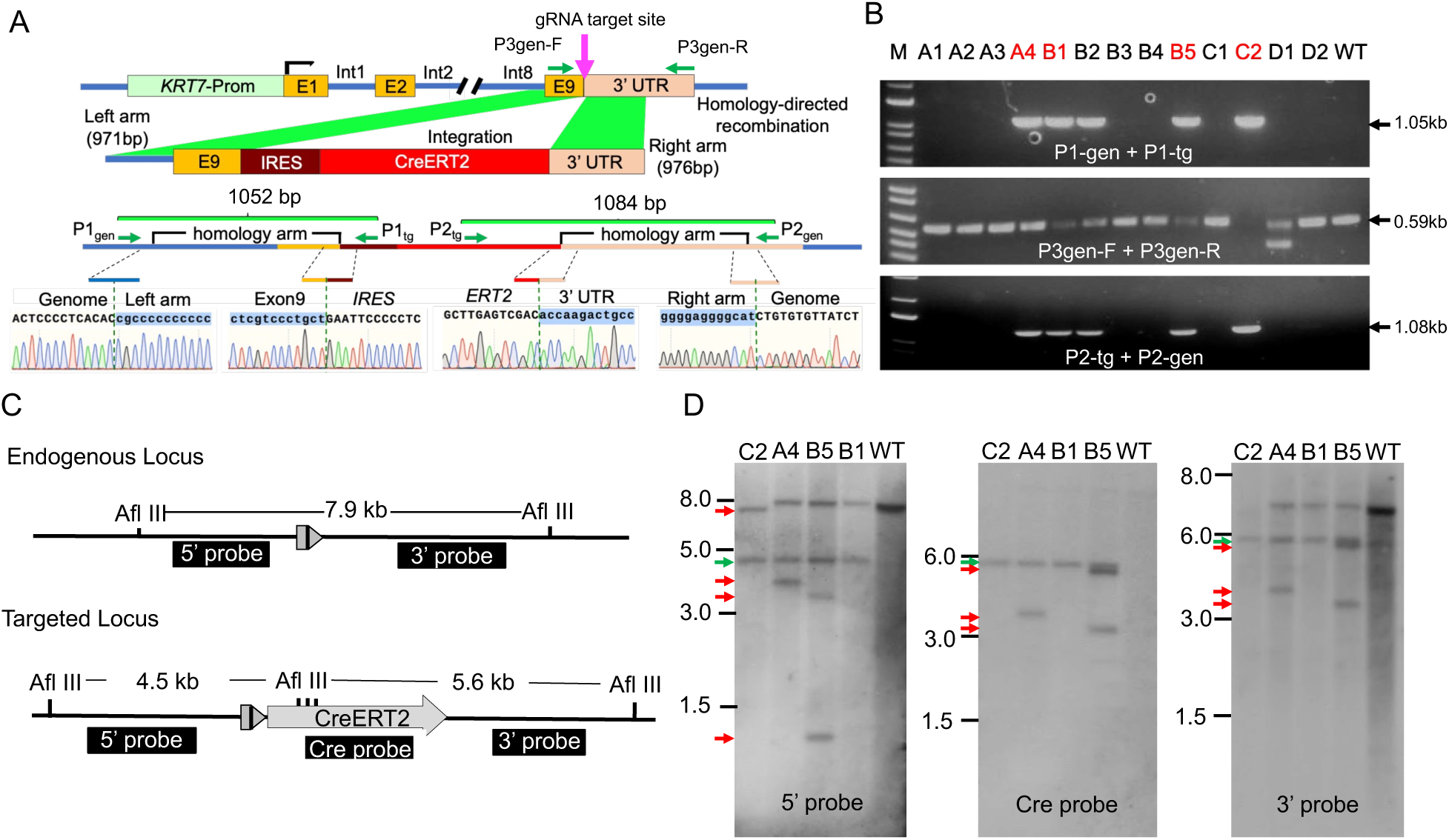
Generation of the *KRT7*^CreER^ ferret line targeting glandular and ductal epithelial cells. (A) Schematic representation of the wild-type allele, donor construct, and targeted allele. The donor construct contains the *IRES-CreERT2*-polyA cassette. The IRES enables cap-independent translation initiation. The CRISPR sgRNA targets the 3′ UTR of the ferret *KRT7* gene (5′-CACCAGACTGACCATTGTGCCGG-3′). PCR primer locations are indicated. Primer pairs P1gen/P1tg and P2tg/P2gen amplify across the 5′ and 3′ homology arms, respectively, whereas P3gen-F/P3gen-R detect the endogenous allele. (B) PCR genotyping shows the expected 1052-bp (P1gen/P1tg) and 1084-bp (P2tg/P2gen) products in knock-in animals but not in wild-type ferrets. Amplification of the endogenous allele (567 bp) distinguishes heterozygous from homozygous animals. (C) Southern blot strategy for validating correct targeting of the *KRT7-CreERT2* allele. (D) Southern blot analysis of F0 founders. All four live founders exhibited the expected band (green arrows). Three founders also showed additional unexpected bands (red arrows).

Four live F0 founders were analyzed by Southern blotting. All founders displayed the correctly targeted insertion. Founder C2 carried an approximately 500 bp deletion in the non-targeted allele, founder A4 exhibited an additional insertion, and founder B5 harbored two extra insertions (Fig.1C-D). Founder B1 showed only the correctly targeted insertion, with no additional integration events. To determine whether the extra insertions in founders A4 and B5 were randomly located or linked to the targeted allele, F1 progeny from each founder were analyzed by Southern blotting (Fig. 2). F1 kits derived from founders B1 and C2 carried only the correctly targeted insertion, whereas F1 kits from founders A4 and B5 retained additional insertions. Given the limited number of F1 offspring from founders A4 and B5, it remains unclear whether these extra insertions are closely linked to the targeted locus. Accordingly, only founders B1 and C2 were selected as breeders for colony establishment.

**Figure 2.**
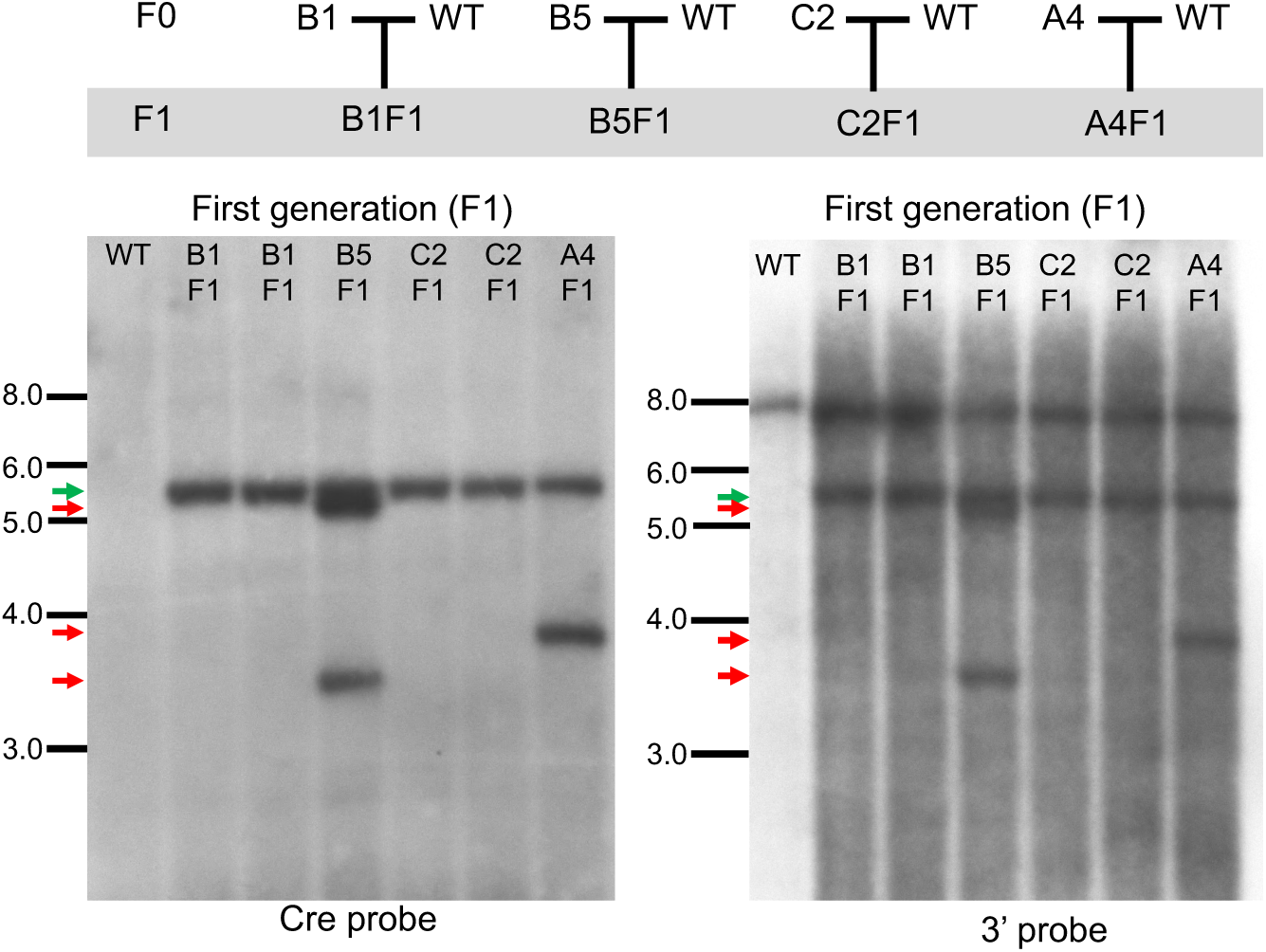
Southern blot analysis of F1 progeny. Southern blotting of genomic DNA from F1 offspring demonstrates inheritance of the targeted *KRT7^CreER^* allele. F1 animals derived from founders A4 and B5 exhibited hybridization patterns identical to their respective F0 founders when probed with Cre-specific and 3′ probes, confirming transmission of the additional insertions. F1 progeny from founder C2 showed the correct insertion pattern without the unexpected band observed in the C2 F0 animal. Green arrows mark expected targeted bands, and red arrows mark unexpected bands.

### Lineage tracing of KRT7-expressing cells in multiple organs under physiological conditions

To enable in vivo lineage tracing of KRT7-expressing cells, *KRT7*^CreER^ founder ferrets were crossed with *ROSA26-LoxP-tdTomato-stop-LoxP-EGFP* (*ROSA^mTmG^*) Cre-reporter ferrets (30). This reporter line contains a dual fluorescent transgene that permanently switches from tdTomato to EGFP expression following tamoxifen-induced activation of CreERT2. Following tamoxifen administration to *KRT7*^CreER^*; ROSA*^mTmG^ animals, EGFP-positive cells reliably marked known sites of KRT7 expression. Consistent with the established epithelial distribution of KRT7, EGFP-positive lineage-labeled cells were found across multiple organ systems (Fig. 3).

**Figure 3.**
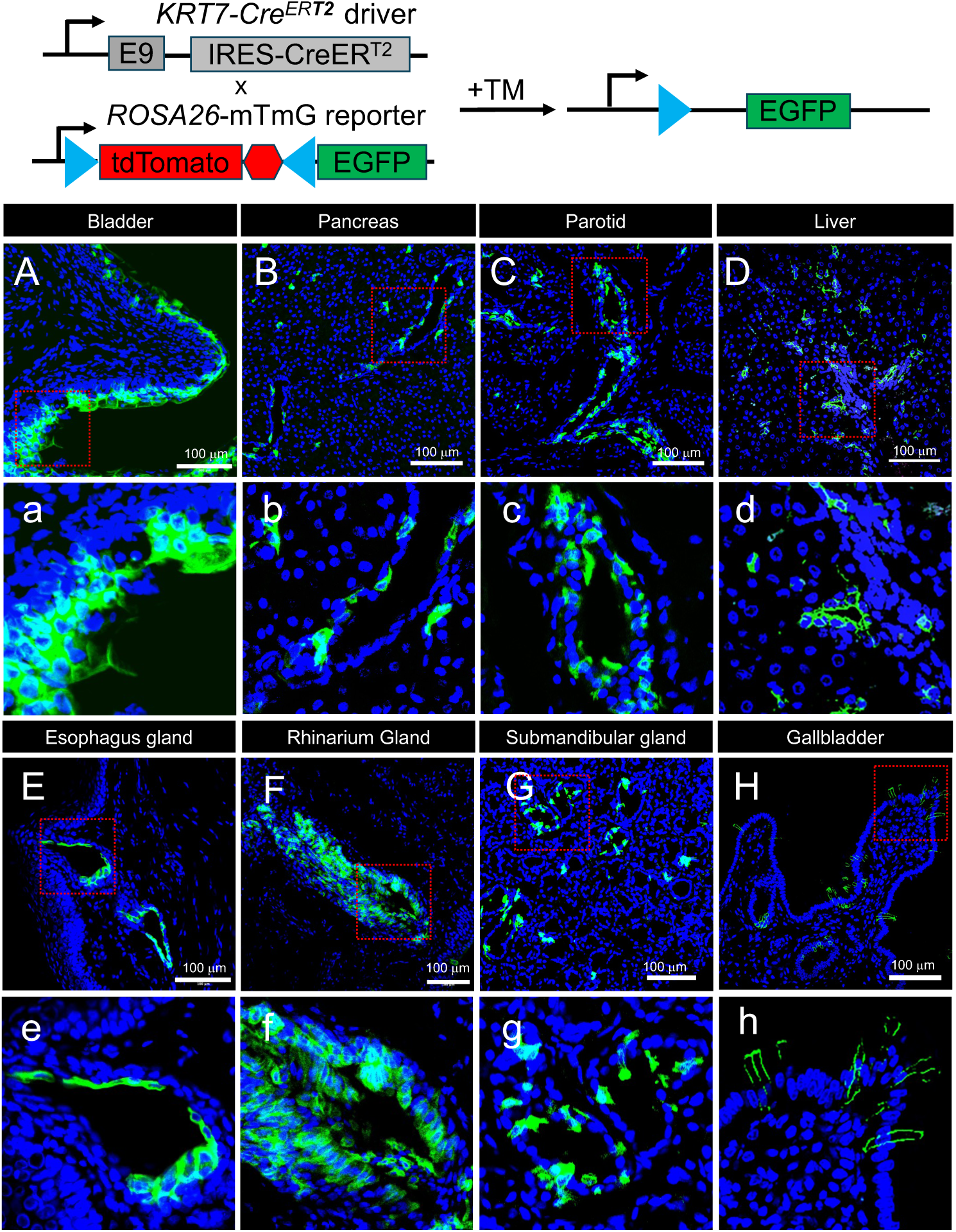
Labeled KRT7⁺ cells are found in multiple organs of *KRT7^Cre^*^ER^*;ROSA*^mTmG^ ferrets. Top panel: schematic diagram of the lineage-tracing method used to visualize KRT7-expressing epithelial cells in vivo. *KRT7^CreER^* ferrets were crossed with the *ROSA26-LoxP-tdTomato-stop-LoxP-EGFP* (*ROSA*^mTmG^) reporter line. In this system, all cells initially express membrane-localized tdTomato; after tamoxifen-induced activation of CreERT2, the tdTomato cassette is excised, resulting in permanent EGFP expression in KRT7⁺ cells and their descendants. Adult *KRT7*^CreER^*;ROSA*^mTmG^ ferrets received intraperitoneal tamoxifen injections for five days in a row. Bottom panel: tissues were collected seven days after the last dose to evaluate EGFP labeling. (A–H) Tamoxifen induction led to strong EGFP expression across multiple organs, revealing widespread distribution of KRT7⁺ epithelial cells. EGFP⁺ cells appeared in glandular and ductal epithelial structures in the (A, a) bladder, (B, b) pancreas, (C, c) parotid gland, (D, d) liver, (E, e) esophagus gland, (F, f) Rhinarium gland, (G, g) submandibular gland, and (H, h) gallbladder. Images from three different animals are shown. Scale bar, 100 μm. TAM, tamoxifen.

Strong labeling appeared in the urothelium of the bladder, the glandular epithelium of the esophagus, and various glandular tissues, including the major salivary glands (parotid and submandibular). Within the pancreas, EGFP-positive cells were prominently detected in terminal and intercalated pancreatic ductal epithelial cells, demonstrating effective recombination in this area. In addition to these classical KRT7-expressing epithelia, scattered EGFP-positive cells were also observed in the epithelial lining of the liver and gallbladder, indicating that subsets of cholangiocytes and biliary epithelial cells express KRT7 at levels sufficient for recombination. The labeled cells were also found in rhinarium glands. The distribution of EGFP-positive cells across these tissues supports the utility of the *KRT7*^CreER^ line for tracking KRT7-positive epithelial lineages across multiple ferret tissues.

### Lineage tracing of KRT7-expressing cells in the salivary gland

To validate that EGFP⁺ cells originated from KRT7-expressing cells, we performed co-immunostaining for KRT7 and EGFP in the salivary gland, where EGFP lineage tracing robustly labeled ductal structures. KRT7 signal colocalized with EGFP in these ducts (Fig. 4), confirming that EGFP⁺ cells correspond to KRT7-expressing ductal epithelial cells. As observed with many Cre-drivers, not all lineage⁺ cells were labeled by EGFP (Fig. 4, arrows). Whether this reflects transcriptional heterogeneity in KRT7 expression within salivary ductal cells and/or inefficient recombination at the tamoxifen dose used remains to be determined.

**Figure 4.**
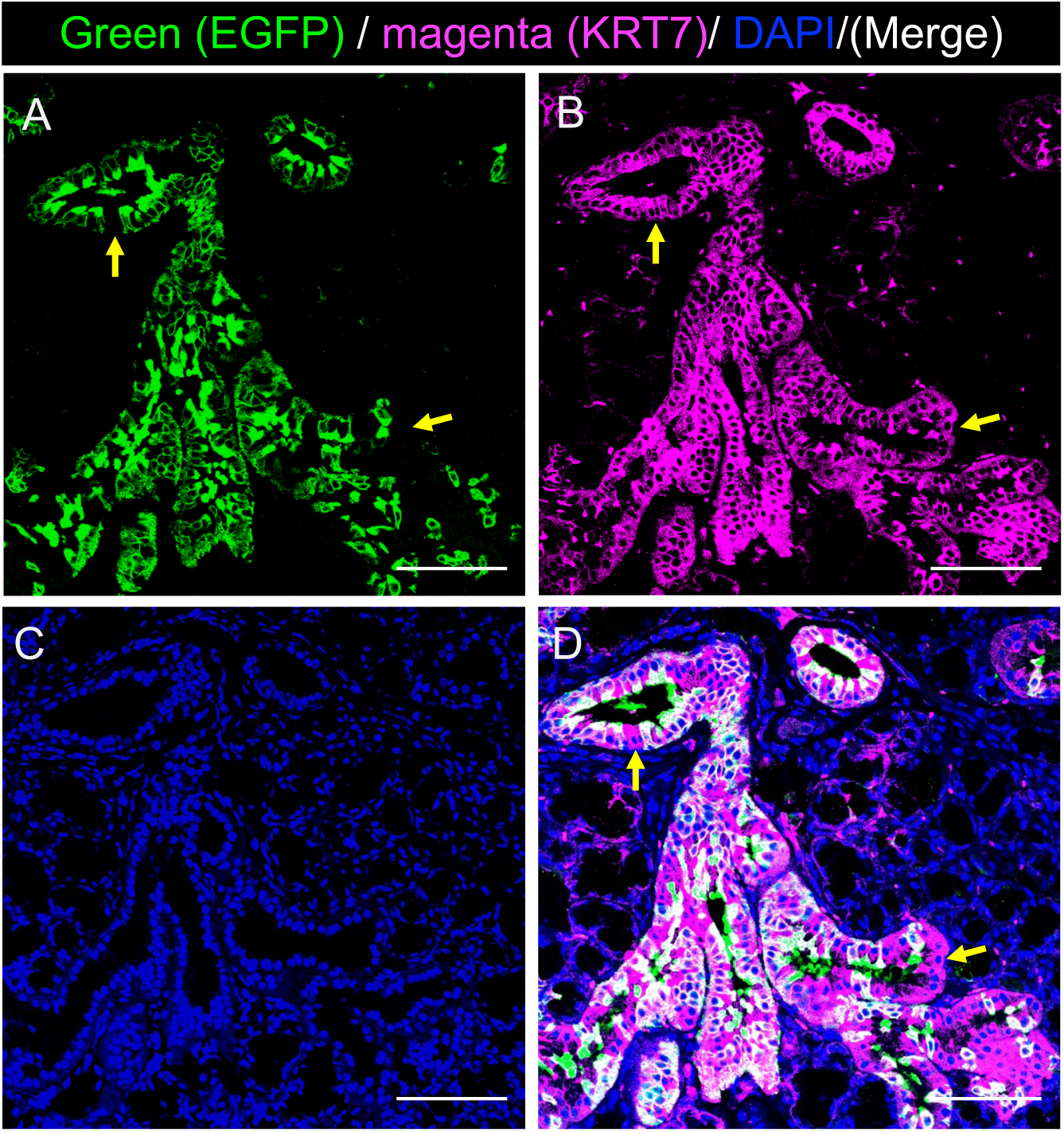
Validation of EGFP labeling in KRT7-expressing salivary duct cells. (A–B) KRT7 signal colocalizes with EGFP in salivary ductal structures, confirming that EGFP⁺ cells represent KRT7-expressing duct cells. (C–D) A subset of KRT7⁺ duct cells lacks EGFP labeling, indicating incomplete lineage tracing. This may reflect suboptimal tamoxifen induction efficiency or cellular heterogeneity within the salivary gland ductal compartment, with only a subset of duct cells being traceable under the current conditions.

### KRT7 lineage tracing of salivary gland ductal cells during postnatal development

Recent single-cell RNA-seq studies have demonstrated that the mouse submandibular gland contains multiple transcriptionally distinct epithelial populations, including diverse ductal, acinar, basal, and myoepithelial cell states (31, 32). Complementary lineage tracing studies show that multipotent Trp63^+^ stem cells give rise to differentiated Krt7^+^ luminal ductal cells (31). To examine ductal cell heterogeneity and turnover within the submandibular and parotid gland, ferrets were pulsed with tamoxifen at one month of age and then followed until six months of age. Because KRT7 has been reported in salivary ductal epithelial populations, including subsets with ionocyte-like transcriptional features, we examined the overlap between KRT7 lineage-traced cells and BSND, a canonical molecular marker of ionocytes previously described in pulmonary gland ducts (33, 34). Immediately after tamoxifen treatment, nearly all luminal ductal cells in the parotid and submandibular glands were lineage labeled by *KRT7*^CreER^ (Fig. 5A-D). By contrast, following the 6-month chase period, the majority of EGFP⁺ cells lacked BSND expression, with only rare cells retaining colocalization (Fig. 5E-H). Collectively, these findings suggest that BSND-negative KRT7-lineage luminal ductal cells are long-lived, whereas BSND-positive ionocyte-like ductal cells undergo more rapid turnover. Thus, KRT7⁺BSND⁺ ionocyte-like ductal cells appear to be replenished over time by a lineage-negative progenitor (e.g., multipotent TP63^+^ stem cells).

**Figure 5.**
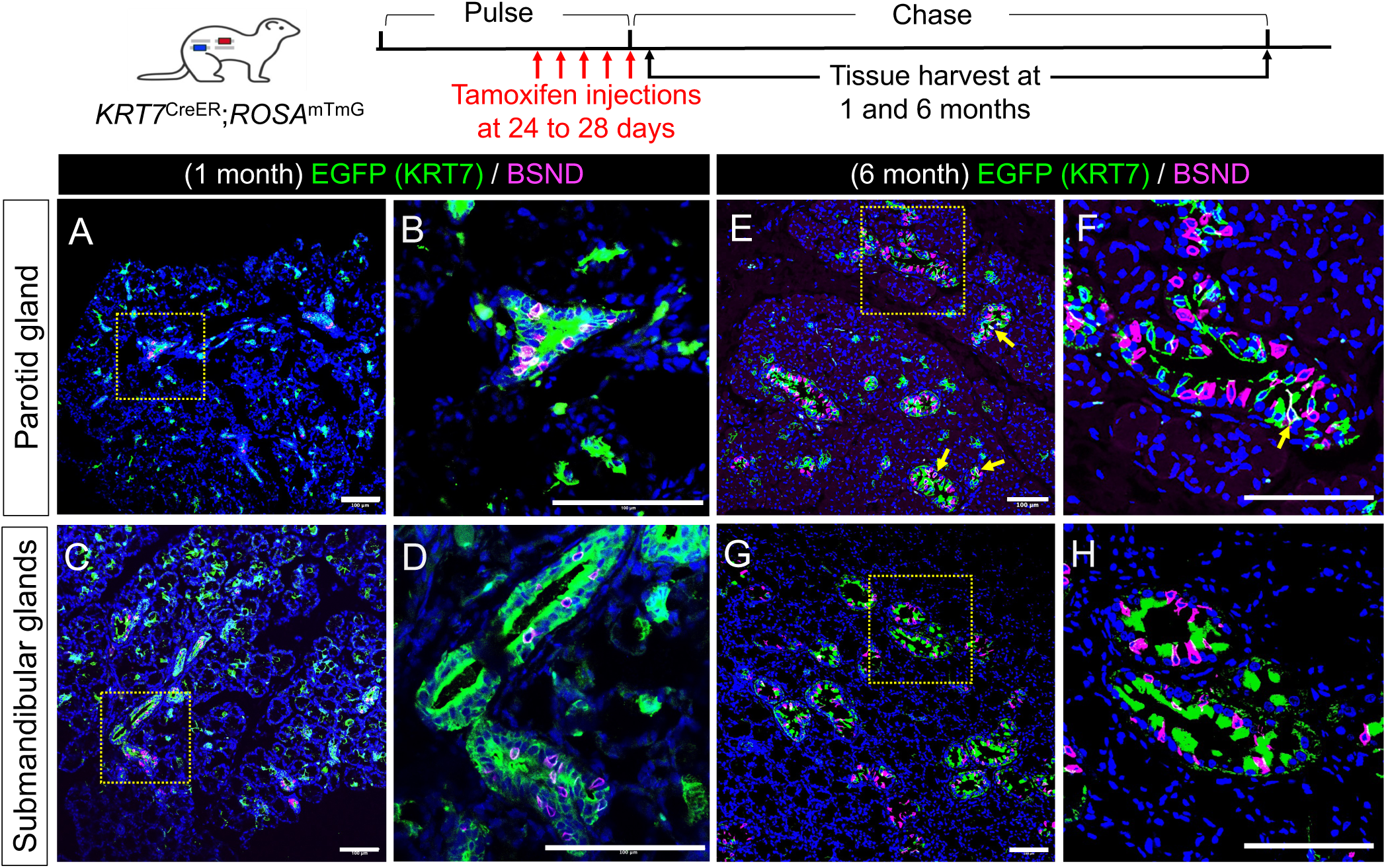
Ductal labeling in salivary glands. Top panel: schematic diagram of the push and chase lineage-tracing method used to visualize KRT7-expressing epithelial cells in the salivary gland in vivo. Bottom panel: (A–D) Lineage-traced EGFP⁺ cells in the ductal compartment of the parotid gland and submandibular gland at 1 month old. Most EGFP⁺ lineage-traced cells were BSND-negative, consistent with a non-ionocyte ductal identity. EGFP co-stained with BSND. (E–F) Lineage-traced EGFP⁺ cells in the ductal compartment of the parotid submandibular gland 5 months after pulsing at 1 month old. The majority of EGFP⁺ cells were BSND-negative, indicating that they predominantly represent non-ionocyte ductal cells. Rare EGFP⁺ cells co-stained with BSND (yellow arrows). KRT7 (EGFP) is a marker of ductal epithelial cells in the salivary gland, whereas BSND (magenta) is a specific marker of ionocytes (specialized ion-transporting cells).

### Fate mapping KRT7-lineage cells in the pancreas under pathological conditions

KRT7 is expressed in pancreatic ductal epithelium (8). During pancreatitis and pancreatic cancer, epithelial remodeling characterized by acinar-to-ductal metaplasia and ductal expansion gives rise to duct-like cell states that may increase KRT7 expression (35). To investigate the behavior of pancreatic KRT7⁺ cells in the context of disease, we conducted lineage-tracing studies in cystic fibrosis (CF) ferrets harboring homozygous *CFTR-*G551D mutations (16, 36). *CFTR*^G551D/G551D^ ferrets are responsive to the CFTR potentiator Ivacaftor (VX-770), a small molecule that binds to the mutant CFTR protein and restores channel opening (16). Thus, administration of VX-770 in utero and postnatally to *CFTR*^G551D/G551D^ ferrets can be used to control the timing of disease onset and progression. CF ferrets develop rapidly progressive pancreatitis within the first two months of life, characterized by fibrosis, inflammation, and complete loss of acinar cells (14–16, 37). Pancreatic remodeling also affects glycemic status early in life through islet loss and gives rise to cystic fibrosis-related diabetes later in life (15, 17, 18). Taken together, pancreatic remodeling in the CF ferret captures multicellular pathological processes that are well-suited to evaluate changes in ductal cell behavior.

*KRT7*^CreER^; *ROSA*^mTmG^ wild-type (WT) ferrets and *KRT7*^CreER^; *ROSA*^mTmG^; *CFTR*^G551D/G551D^ CF ferrets received five daily doses of tamoxifen (15 mg/kg) at one month of age. To preserve pancreatic function, CF ferrets were maintained on VX-770 in utero and postnatally until 10 days after the tamoxifen induction. VX-770 was then discontinued to trigger CF pancreatitis, and animals were euthanized 10 days later. In the wild-type (WT) pancreas, EGFP⁺ cells were readily detected in intercalated and intralobar ducts (Fig. 6A), terminal ductal (centroacinar) cells (Fig. 6A, B), and main ducts (data not shown). In the setting of early CF pancreatitis, EGFP labeling persisted in ductal cells within dilated terminal ducts and enlarged intercalated and intralobar ducts, demonstrating that KRT7-lineage cell fate can be tracked during CF pancreatic remodeling (Fig. 6C–G).

**Figure 6.**
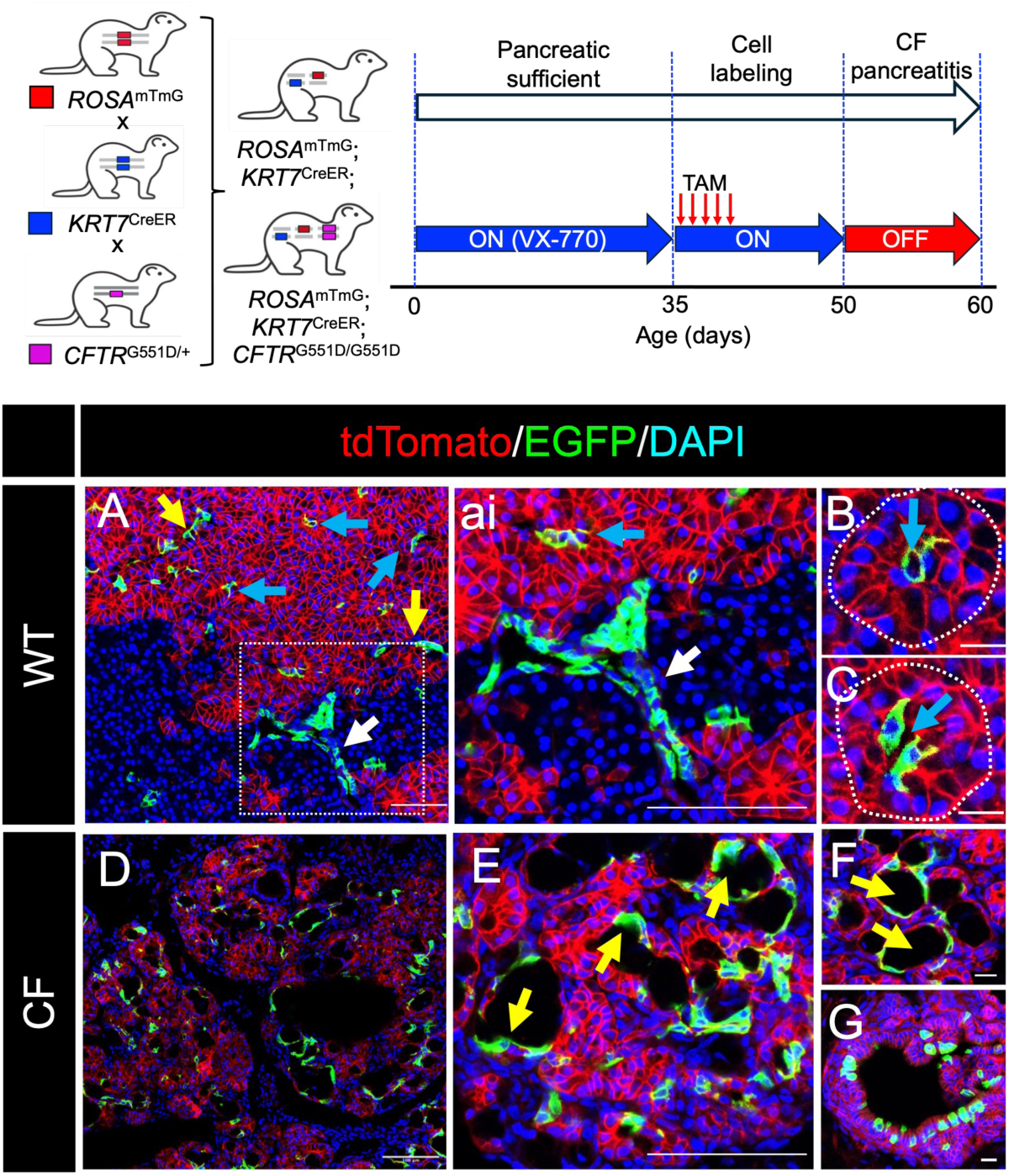
*KRT7*^CreER^;*ROSA*^mTmG^ lineage tracing in WT and CF ferret pancreas. Top panel: Breeding scheme for generating *KRT7^CreER^;ROSA*^mTmG^*;CFTR*^G551D/G551D^ triple-transgenic ferrets and timeline of tamoxifen administration used to induce CreERT2-mediated recombination. **Bottom panel:** KRT7-derived lineage tracing in WT and CF pancreas. (A) In WT pancreas, EGFP lineage-labeled cells are present in multiple ductal epithelial compartments. EGFP⁺ intercalated duct cells are indicated by yellow arrows, terminal ductal/centroacinar cells (CACs) by blue arrows, and intralobar ductal cells by white arrows. (ai) Higher-magnification view of the boxed region in (A), highlighting the morphology and spatial organization of EGFP⁺ lineage-labeled ductal epithelial cells. (B, C) White dotted outlines demarcate acinar units, illustrating the localization of EGFP⁺ CACs at the junction between ducts and acini. (D) In the CF pancreas, EGFP lineage-labeled cells are retained despite chronic inflammatory and obstructive pathology, demonstrating persistence of KRT7-derived ductal lineages under disease conditions. (E) Swollen intercalated ducts (yellow arrows) characteristic of CF pancreatitis. (F) Enlarged terminal ducts containing EGFP⁺ lineage-labeled epithelial cells. (G) EGFP⁺ cells present in intralobar ducts of CF pancreas, indicating maintenance of KRT7-derived ductal lineages across multiple ductal compartments in the diseased state.

To further characterize lineage-labeled cells (EGFP⁺), we performed co-immunostaining for SOX9, a transcription factor that plays a key role in maintaining pancreatic ductal cell identity and homeostasis (38, 39). In the pancreas, SOX9 and KRT7 expression are broadly associated with ductal epithelial lineages, extending from centroacinar/terminal duct cells to larger duct compartments, where they overlap with CFTR expression (40–44). Lineage tracing of *KRT7*^CreER^; *ROSA*^mTmG^ ferrets revealed distinct differences in the organization of SOX9⁺ centroacinar cells (CACs) and intercalated ducts between WT and CF ferret pancreas. In WT tissue, lineage labeling (i.e., EGFP^+^) was confined to SOX9⁺ CACs located within acinar units and to SOX9⁺ intercalated ducts with preserved morphology (Fig. 7A–D), consistent with stable maintenance of SOX9⁺ ductal and centroacinar compartments under non-diseased conditions. In contrast, the CF pancreas exhibited prominent alterations within the SOX9⁺ ductal lineage. Lineage labeling highlighted markedly dilated SOX9⁺ intercalated ducts (Fig. 7E, eii), consistent with ductal structural abnormalities associated with CF pathology. In addition, hyperplastic EGFP^+^SOX9⁺ pod-like structures were observed in terminal ductal regions (Fig. 7E, ei, F), indicating localized remodeling of ductal architecture. Collectively, these findings indicate that the initial stages of CF pancreatitis are associated with abnormal expansion of KRT7 lineages marking intercalated and terminal ductal cells.

**Figure 7.**
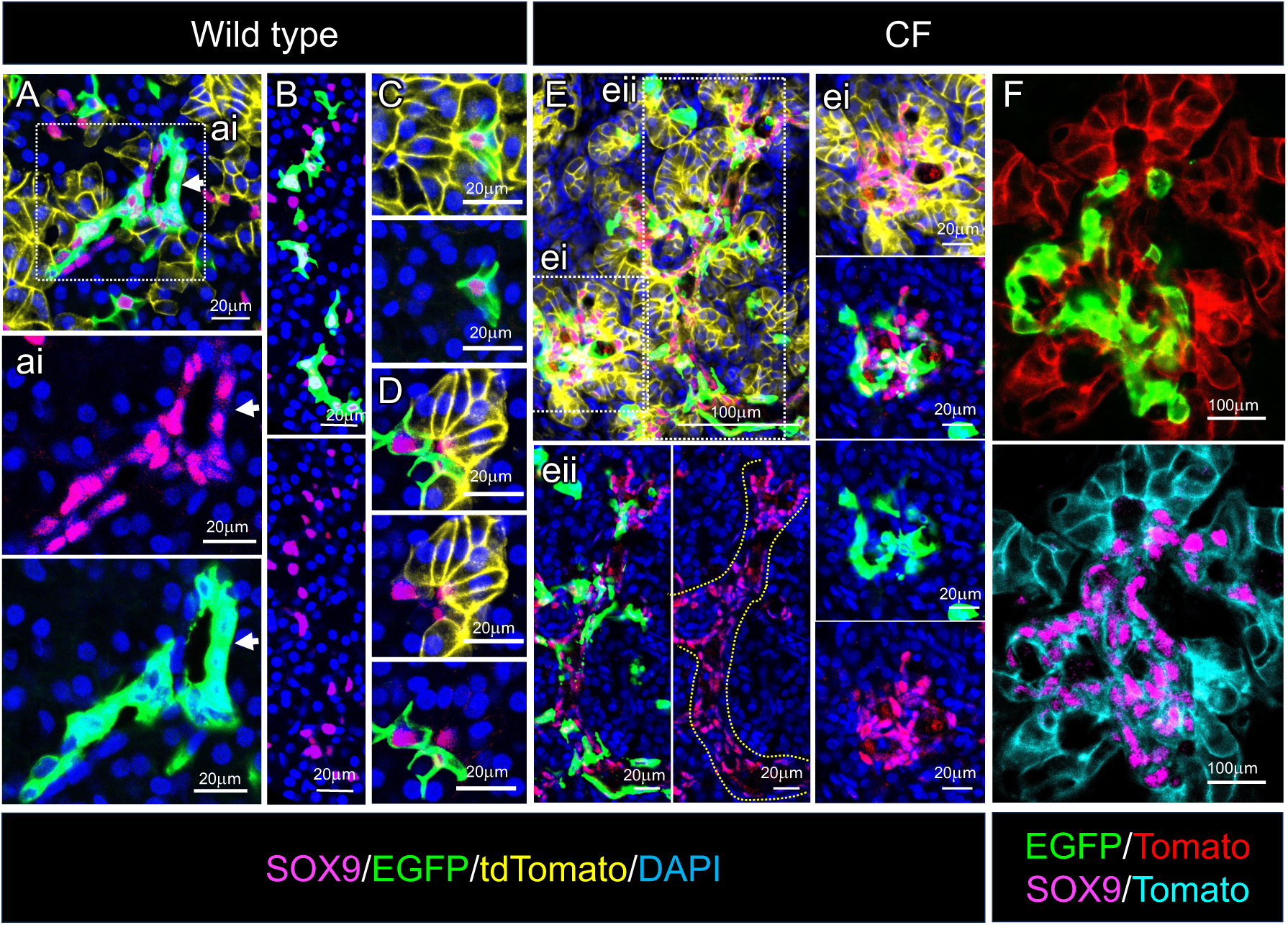
Lineage labeling of SOX9⁺ centroacinar cells (CACs) and intercalated ducts in *KRT7*^CreER^; *ROSA*^mTmG^ wild-type and CF ferrets. (A–B) In WT pancreas, lineage tracing labels SOX9⁺ CACs located within acinar units. (C–D) SOX9⁺ intercalated ducts are similarly labeled in WT tissue. (E) In the CF pancreas, lineage labeling highlights (ei) markedly swollen SOX9⁺ intercalated ducts and (eii) SOX9⁺ pod-like structures emerging in terminal ductal regions. In (eii), the white dotted outline delineates the intercalated duct. (F) Enlarged SOX9⁺ pod structures containing EGFP⁺/SOX9⁺ cells, consistent with expansion of intercalated duct-derived cells.

### KRT7 lineage ductal cells acquire an acinar trait in the injured CF state

During pancreatitis or early CF-associated pancreatic remodeling, acinar, centroacinar, and ductal cells can acquire programs that drive epithelial plasticity and alter subsequent fate decisions and lineage identity (15, 45, 46). To assess the extent to which this might occur in the CF pancreas, we localized a well-known acinar-specific gene (CELA3B) together with EGFP expression from KTR7 lineage-traced cells. In the wild-type (WT) *KRT7*^CreER^; *ROSA*^mTmG^ pancreas, immunohistochemistry and immunofluorescent smFISH analyses revealed no *EGFP*⁺ cells co-expressing *CELA3B*, indicating that acinar cells were not labeled under physiological conditions. Instead, EGFP labeling was observed in centroacinar cells as well as intercalated and intralobular ducts (Fig. 8A–C). In contrast, analysis of the cystic fibrosis (CF) pancreas revealed EGFP⁺ cells that co-expressed CELA3B (Fig. 8D–F). These *EGFP⁺/CELA3B*⁺ cells were detected by smFISH, indicating that acinar cells undergo acinar-to-ductal metaplasia (ADM) in the CF pancreas and can be lineage-labeled by the *KRT7-CreERT2* system under disease conditions. To further investigate *KRT7-CreERT2*–mediated labeling in acinar-derived ADM contexts, in vitro models of pancreatic ADM were analyzed. Cystic organoids generated from cultured single acinar cells expressed tdTomato in the absence of 4-hydroxytamoxifen (4-OHT) (Fig. 8G–H). Following 4-OHT treatment, these organoids expressed EGFP (Fig. 8I), demonstrating Cre-dependent reporter activation during acinar-derived duct-like remodeling in vitro. EGFP⁺/CELA3B^+^ cells may also come from ditransdifferentiated duct cells caused by the mutation of CFTR.

**Figure 8.**
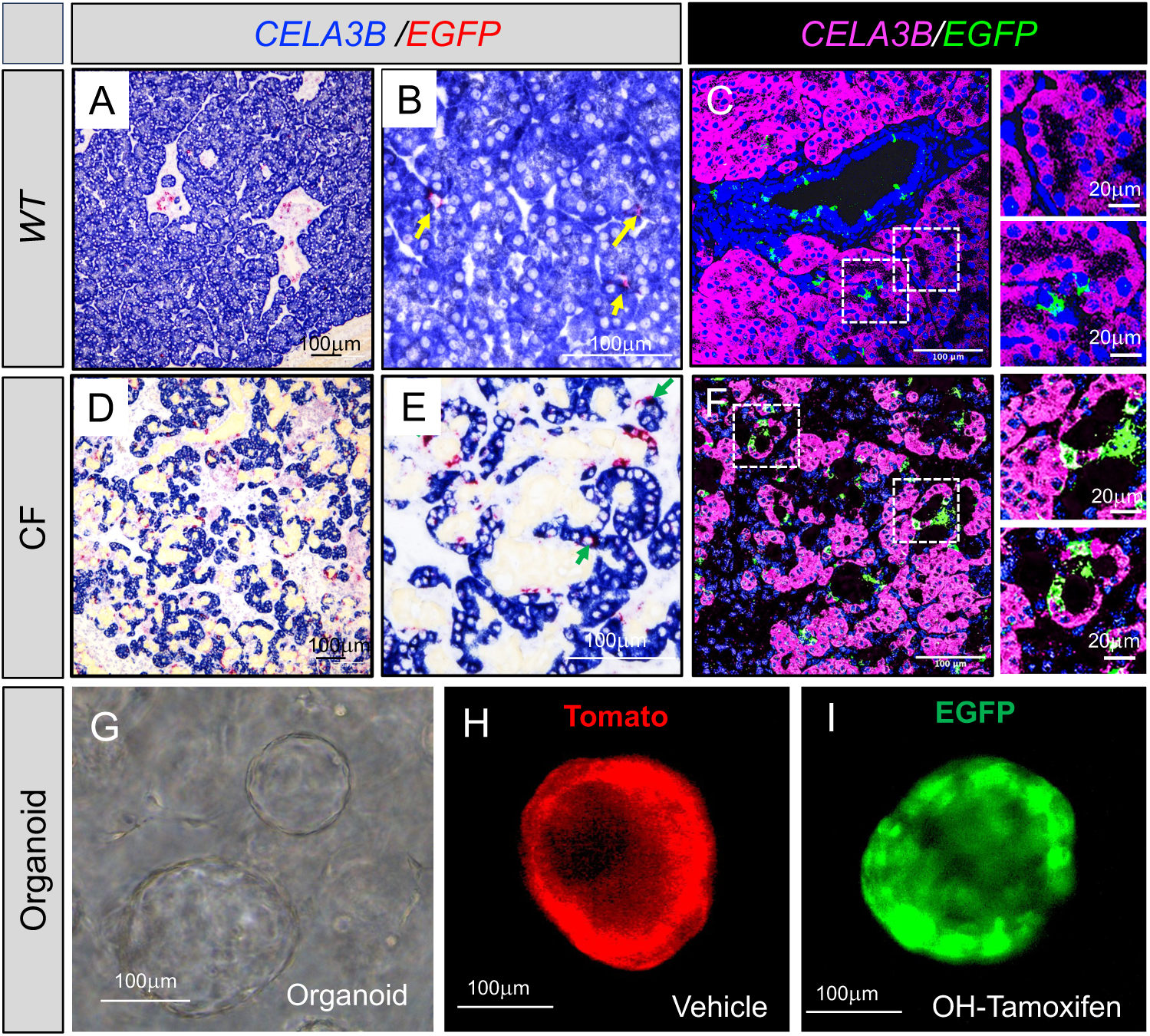
Acinar cells are not labeled in the WT pancreas but are labeled in the CF pancreas. (A– B, D–E) Immunohistochemistry and (C, F) immunofluorescent smFISH images detecting *EGFP* and *CELA3B* mRNA. (A–C) In the WT pancreas, no EGFP⁺ cells co-express *CELA3B*, indicating that acinar cells are not labeled under physiological conditions. EGFP⁺ cells are observed in the centroacinar and ductal compartments. (D–F) In the CF pancreas, some *EGFP⁺* cells co-express *CELA3B,* consistent with labeling of acinar cells undergoing acinar-to-ductal metaplasia. (G–I) In vitro ADM model derived from cultured single acinar cells. (G) Cystic organoid formed from acinar cells. (H) In the absence of 4-OHT, organoids express tdTomato. (I) Following 4-OHT treatment, organoids express EGFP, demonstrating Cre-dependent reporter activation.

## Discussion

We used the Easi-CRISPR method to generate the *KRT7*^CreER^ ferrets. Easi-CRISPR relies on pronuclear injection of a long single-stranded DNA donor (lssDNA) to facilitate precise genome editing (47). Mouse studies have shown that lssDNA reduces random genomic integration compared with double-stranded DNA donors (47). However, our findings demonstrate that random insertions occurred in 50% of founder ferrets generated by Easi-CRISPR, underscoring the need for rigorous screening of F1 offspring using Southern blotting or other whole-genome sequencing–based approaches.

To determine KRT7 expression in various organs in ferrets, we performed lineage tracing in this *KRT7*^CreER^ model. We identified cells predominantly localized to simple epithelia across multiple organs, including the pancreas, salivary gland, liver, esophageal gland, and gallbladder. These distribution and expression patterns are consistent with the observation in humans. (48–53). While KRT7 is classically associated with simple epithelia, human studies have noted a degree of expression promiscuity, with KRT7 also detected in tissues containing stratified epithelia, such as the bladder epithelium. (54). We observed a similar pattern in ferrets. We also observed that KRT7 is expressed in the rhinarium gland of the ferret, a finding not previously reported. These findings support both the evolutionary conservation of KRT7 expression and the tissue specificity of the KRT7-IRES-CreER-based lineage-tracing strategy in ferrets, validating its utility for identifying and tracking simple epithelial cell populations across organs.

Recent transcriptomic studies have identified a specialized subpopulation of salivary gland duct cells, called salivary ionocytes, that express FGF10 and ASCL3 (55, 56), but ionocytes in pups younger than 5 days are FGF10-negative (56). Comparative analyses show that ionocytes are conserved across organs and share a characteristic gene expression program, including KRT7(57). However, whether the KR7 is expressed in neonatal ionocytes or mature ionocytes remains unknown. To this end, we performed a pulse-and-chase lineage-tracing experiment in these new models. KRT7 lineage tracing uniformly labeled most luminal ductal cells in 1-month-old ferrets, including BSND^+^ ionocyte-like cells. Five months later, the persisting lineage-labeled population was predominantly BSND-negative, suggesting more rapid turnover of the ionocyte population and replacement by lineage-negative progenitors. In contrast to our findings in ferrets, lineage tracing of mouse salivary gland ionocytes using *Fgf10*^CreER^ revealed persistent labeling for up to 8 months (56), suggesting that ionocyte turnover may differ across species. Notably, scRNA-seq has identified Krt7 as part of the conserved ionocyte transcriptional signature in mouse salivary gland (56) and more broadly ionocytes from multiple human organs (57). To our knowledge, Krt7-based fate mapping has not been previously applied to mouse salivary gland epithelia. Thus, the *KRT7*^CreER^ ferret provides a new avenue to study luminal ductal and ionocyte-associated lineage dynamics and reveals previously unappreciated cellular heterogeneity within ferret salivary ducts.

The pancreas provided the clearest demonstration of the translational value of the *KRT7*^CreER^ ferret model. Under homeostatic conditions, KRT7-lineage labeling localized to exocrine ductal compartments, including centroacinar and intercalated duct cells, supporting prior studies in human and rodent pancreas that identify KRT7 as a marker of ductal epithelium rather than mature acinar or endocrine cells (8, 58, 59). In the setting of CF pancreatic injury, KRT7-lineage cells underwent prominent remodeling characterized by apparent pod-like expansion of SOX9-positive terminal ductal structures into enlarging intercalated ducts. Because SOX9 regulates pancreatic ductal identity, progenitor responses, and epithelial repair after injury (38, 39, 42), these changes may reflect an injury-responsive program that includes ADM, as also observed in caerulein-induced and pancreatic duct ligation models of pancreatitis (45, 60, 61). There is another explanation for KRT7^+^/CELA3B^+^ cells. In ferret CF pancreas, loss of CFTR potentiation unmasks injury-associated plasticity in fate-mapped SOX9^+^/KRT7^+^ centroacinar/ductal descendants. Unlike WT, these EGFP+ descendants can acquire an acinar-enzyme program, including CELA3B, consistent with CAC/ductal-to-acinar lineage plasticity or attempted exocrine regeneration after CFTR-dependent pancreatic stress, rather than baseline CAC expression of *CELA3B.* These findings also support emerging evidence that small ductal and centroacinar populations are dynamic participants in pancreatic regeneration and maintain clonogenic multipotent properties in culture (41, 46).

This study has several limitations. Recombination was incomplete in certain organs, such as the pancreas, and therefore lineage labeling likely underestimates the full size of KRT7-positive compartments. Our pancreatic studies focused on an early injury window; later stages may reveal additional lineage transitions or regenerative outcomes. Finally, lineage tracing identifies cell history but does not, by itself, define function; future studies combining this model with transcriptomic or cell-ablation approaches will be informative.

In summary, *KRT7^C^*^reER^ ferrets provide a new tool for studying epithelial lineage dynamics in a clinically relevant species. The model enables inducible fate mapping of ductal and glandular epithelial populations across multiple organs under both homeostatic and disease conditions, providing a versatile tool for investigating lineage stability, cellular plasticity, and tissue remodeling. These capabilities should accelerate studies of exocrine tissue repair, epithelial plasticity, and disease mechanisms that are not adequately captured in traditional rodent systems.

## Methods

### Sex as a biological variable

Sex was not considered a biological variable, and both female and male ferrets were used in this experiment.

### Animals

The Institutional Animal Care and Use Committee (IACUC) of the University of Iowa and the University of Alabama at Birmingham approved this study. *ROSA*^mTmG^ and CFTR^G551D^ ferrets were previously described (30, 36). Triple transgenic ferrets (*KRT7*^CreER^; *ROSA*^mTmG^; CFTR^G551D^) were generated by breeding *KRT7*^CreER^; *ROSA*^mTmG^ ferrets with CFTR^G551D^ ferrets. Intraperitoneal (IP) injections of tamoxifen (Tam), solubilized by sonication in 100% sunflower seed oil (100 mg/kg body weight, 20 mg/mL stock solution, #T5648; Millipore Sigma, St. Louis, MO), were administered daily for five consecutive days at the specified ages (62). CFTR^G551D/G551D^ CF ferrets were reared in utero and postnatal VX-770 as previously described (16).

### Generation of the *KRT7*^CreER^ driver ferret line

The founder KRT7-IRES-CreERT2 driver ferret was generated using CRISPR-mediated Homologous Recombination (HR) in zygotes to insert an IRES-CreERT2 cassette downstream to the stop codon in the 3’UTR of *KRT7* (NW_025422368.1; 3657953..3671995, complement). The IRES-CreERT2 cassette was flanked by ∼1kb homology arms and cloned into a plasmid that allowed for single-stranded DNA generation using the Guide-it Long ssDNA Production System v2 from Takarabio Inc (Cat #632666). The single-stranded DNA was gel-isolated and purified using a NucleoSpin Gel and PCR Clean-Up from Macherey-Nagel Inc. (Cat. No. 740609.50). gRNAs to the targeted region were screened by in vitro cleavage assays on G-Block DNA. The ribonucleoprotein, the spCas9 complexed with gRNA, and the long single-stranded DNA donor were microinjected into zygotes. Injected zygotes were cultured in TCM-199 medium supplemented with 10% fetal calf serum (FCS) overnight to the 2-cell stage, then transferred into primiparous pseudo-pregnant jills. The kits were naturally delivered in 42 days (full-term gestation). The gRNA was designed using the online tool CRISPOR (http://crispor.tefor.net). spCas9 protein (from *Streptococcus pyogenes*), gRNA, and PCR primers for genotyping and TaqMan quantitative PCR assays were generated at Integrated DNA Technologies, Inc. (Coralville, IA).

### PCR, Sanger sequencing, and genotyping

The founder animal was identified by PCR-based genotypic analysis, followed by confirmation by Sanger sequencing. Primer sets, P1_gene_/P1_tg_ (P1_gene_: 5’-CATGACTACCCTTCATCACACCCTG-3’; P1_tg_: 5’-CACCGGCCTTATTCCAAGCG-3’) and P2_tg_/P2_gene_ (P2_tg_: 5’-CGGGCTCTACTTCATCGCATTC-3’; P2_gene_: 5’-ATCCCCCCTTCAGACACCCTG-3’) were used to amplify the junction sequences at the 5′ (left) and 3′ (right) of the insert, respectively. The primer pair P3_gene_-F (5’-GCTGTGGTGAACTCCACTGGTG-3’) and P3_gene_-R (5’-TTCTTCCCCAGCCACCTCCAT-3’) amplified the endogenous sequence. Flanking junctional PCR was used to genotype the transgene and WT sequences. Ferret genomic DNA was extracted from tail samples using the Qiagen Gentra Puregene Tissue Kit (Qiagen Sciences, Maryland, USA). PCR amplicons were visualized on standard agarose gels and were confirmed by sequencing after gel extraction.

### Southern blotting

Following restriction enzyme digestion, ferret genomic DNA was resolved on a 1% agarose gel and transferred to a positively charged nylon membrane (GE Healthcare, Chicago, IL) for blotting. Target bands were visualized by autoradiography with radioactive phosphorus-32 (^32^P)-labeled probes, as previously described (63).

### Immunofluorescence analysis

Ferret salivary gland, liver, gallbladder, bladder, esophagus, rhinarium, and pancreas were dissected and fixed overnight in 4% paraformaldehyde (PFA) in phosphate-buffered saline (PBS; pH 7.4). Tissues were embedded in paraffin, sectioned at 4 μm, and processed for immunofluorescence staining. Antigen retrieval was performed by incubation in 10 mM citrate buffer. Sections were washed three times for 5 min each and blocked in 20% donkey serum/PBS for 1–2h at room temperature. Primary antibodies were applied and incubated overnight at 4°C. Sections were then washed three times for 5 min each and incubated with secondary antibodies for 1 h at room temperature. The following antibodies were used: anti-Barttin CLCNK Type Accessory Subunit Beta (BSND) (1:300–1:500; ab196017, Abcam, Waltham, MA), anti-EGFP (1:1000; ab13970 Abcam, Waltham, MA), anti-KRT7 (1:200; MA1-06315, Thermo Fisher Scientific), anti-SOX9 (1:300; AB5535, Millipore Sigma), anti-INS (1:10; P01315-INS, Dako), Alexa Fluor 488 donkey anti-chicken IgG (1:1000; 703-546-155, Jackson ImmunoResearch, West Grove, PA), Alexa Fluor 555 donkey anti-mouse IgG (1:500; A31570, Life Technologies, Carlsbad, CA), Alexa Fluor® 647-AffiniPure F(ab’)2 Fragment Donkey Anti-Guinea Pig IgG (H+L) (706-606-148, Jackson ImmunoResearch), and Alexa Fluor 647 donkey anti-rabbit IgG (1:500; Jackson ImmunoResearch, West Grove, PA). After secondary antibody incubation, samples were washed three times for 5 min each in PBS.

Frozen sections (8-μm) of pancreas embedded in optimal cutting temperature (OCT) compound were air-dried for 30 min at room temperature, then post-fixed in 4% paraformaldehyde (PFA) for 10 min. Tissue sections were permeabilized in 0.2% Triton X-100 in PBS for 30 min, blocked with 5% donkey serum for 2 h, incubated with the primary antibody in diluent buffer (1% donkey serum, 0.03% Triton X-100, and 1 mM CaCl2 in PBS) at 4 °C overnight, and incubated with the Alexa Fluor-labeled secondary antibody at room temperature for 2 h. Slides containing nuclei were stained with DAP or Hoechst 333427 and then mounted with VECTASHIELD Antifade Mounting Medium and imaged on a Zeiss LSM880 confocal microscope or Nikon NSPARC confocal microscopes. All images were processed using FIJI-ImageJ.

### In situ hybridization for pancreatic insulin and glucagon RNA

Pancreata were fixed in 10% neutral-buffered formalin, processed, embedded in paraffin, and sectioned at 10 μm. Sections were baked at 60°C for 1 h, rinsed twice in xylene for 10 min each, incubated twice in 100% ethanol for 10 min each, and air-dried. Slides were processed using the QuantiGene ViewRNA kit (QVT0012; Affymetrix). Slides were incubated in 1× pretreatment buffer (QVT0500; Affymetrix) at 95°C for 10 min, followed by protease digestion at 40°C for 15 min and post-fixation in 10% neutral-buffered formalin for 5 min. Type 1 and type 6 RNA targeting probes were diluted in probe set diluent (QVT0511; Affymetrix) and incubated simultaneously on tissue sections for 3 h at 40°C in a humidified chamber. Slides were washed at room temperature with proprietary Affymetrix wash buffers, then incubated with preamplifier and amplifier oligonucleotides at 40°C for 20 min. Chromogenic signal development was performed by hybridization with alkaline phosphatase–conjugated type 6 detection probes, followed by Fast Blue substrate (QVT0506; Affymetrix) in 0.1 M Tris-maleate buffer. The reaction was terminated with Stop QT buffer (QVT0517; Affymetrix) and repeated using alkaline phosphatase– conjugated type 1 detection probes and Fast Red substrate (QVT0505; Affymetrix) in naphthol buffer. Slides were rinsed in PBS three times for 5 min each and mounted with Vectashield Mounting Medium (Vector Labs, H-1500). All RNA targeting probes were purchased from Affymetrix. Ferret genome sequences for *CELA3B* and *EGFP* were submitted to Affymetrix for probe design and synthesis. Type 6 probes were used to detect ferret insulin mRNA, and type 1 probes were used to detect ferret glucagon mRNA.

## Data availability

All data supporting the findings of this study are available within the article and its Supplementary Information files.

## Author contributions

YY, ZY, XS, and JFE conceptualized the study. YY, XS, and JFE wrote the manuscript. YY, YS, PW, YW, ZH, XL, and ZY performed the experiments. YY, YS, YF, ZY, XS, and JFF interpreted the data. XS, YY, and JFF critically revised the manuscript for important intellectual content. All authors reviewed and edited the manuscript. YY and YS are co-first authors, having contributed equally to the project’s development and the manuscript’s preparation. The order of authorship reflects the relative contributions to the work.

